# Scopolamine and medial frontal stimulus-processing during interval timing

**DOI:** 10.1101/598862

**Authors:** Qiang Zhang, Dennis Jung, Travis Larson, Youngcho Kim, Nandakumar S. Narayanan

## Abstract

Neurodegenerative diseases such as Parkinson’s disease (PD), dementia with Lewy Bodies (DLB), and Alzheimer’s disease (AD) involve loss of cholinergic neurons in the basal forebrain. Here, we investigate how cholinergic dysfunction impacts the frontal cortex during interval timing, a process that can be impaired in PD and AD patients. Interval timing requires participants to estimate an interval of several seconds by making a motor response, and depends on the medial frontal cortex (MFC), which is richly innervated by basal forebrain cholinergic projections. Past work has shown that scopolamine, a muscarinic cholinergic receptor antagonist, reliably impairs interval timing. We tested the hypothesis that scopolamine would attenuate time-related ramping, a key form of temporal processing in the MFC. We recorded neuronal ensembles from 8 mice during performance of a 12-s fixed-interval timing task, which was impaired by the administration of scopolamine. Consistent with past work, scopolamine impaired timing. To our surprise, we found that time-related ramping was unchanged, but stimulus-related activity was enhanced in the MFC. Principal component analyses revealed no consistent changes in time-related ramping components, but did reveal changes in higher components. Taken together, these data indicate that scopolamine changes stimulus-processing rather than temporal processing in the MFC. These data could help understand how cholinergic dysfunction affects cortical circuits in diseases such as PD, DLB, and AD.

**Highlights:**

- The cholinergic muscarinic inhibitor scopolamine impairs interval timing behavior.
- Scopolamine does not change time-related ramping activity in the medial frontal cortex.
- Medial prefrontal stimulus-related modulation increased

## Introduction

Cholinergic dysfunction is a major feature of Alzheimer’s disease (AD), Parkinson’s disease (PD), and dementia with Lewy bodies (DLB) [1-5]. In particular, cholinergic neurons are located in the basal forebrain, which suffers marked neurodegeneration in AD, PD, and DLB [1, 6-9]. Furthermore, drugs that block acetylcholine breakdown, such as donepezil and rivastigmine, can improve cognitive function in PD, DLB, and AD. Basal forebrain cholinergic neurons project broadly to the cortex; however, it is unknown how cholinergic dysfunction affects cortical circuits.

One cognitive process that depends on cholinergic circuits and is consistently impaired in AD and PD is interval timing, which requires subjects to estimate an interval of several seconds by making a motor response [10-13]. Interval timing is ideal for investigating cortical cholinergic function because 1) it depends on the medial frontal cortex (MFC), which is disrupted in AD, PD, and DLB [14-18], 2) it is highly conserved across mammalian species and thus can be readily investigated in rodent models [19], and 3) interval timing in rodents is reliably impaired when they are given scopolamine, a cholinergic inhibitor of M1 muscarinic receptors [20, 21]. Our recent work has indicated that a key form of temporal processing in the rodent MFC is time-related ramping activity; in other words, monotonic increases or decreases in firing rate across a temporal interval [16, 22, 23]. These data lead to the specific hypothesis that scopolamine impairs time-related ramping by MFC neurons.

We tested this hypothesis by recording neuronal ensembles from mice performing a 12-s fixed-interval-timing task, and administering intraperitoneal saline or scopolamine. We found that while scopolamine impaired interval timing, it did not affect time-related ramping in the MFC. Surprisingly, it increased stimulus-related processing in this brain structure. We interpret these data in the context of cholinergic functions and circuits relevant for AD, PD, and DLB.

### Experimental Procedures

#### Mice

This study used 8 wild-type C57/BL6J male mice purchased from Jackson Laboratories (000664) at 3 months of age. Mice consumed 1–1.5 g of sucrose pellets during each behavioral session, and additional food was provided 1–2 hr after each behavioral session in the home cage. Single housing and a 12-hr light/dark cycle were used; all experiments took place during the light cycle. Mice were maintained at 80-85% of their baseline body weight during the course of these experiments for motivation. All procedures were approved by the Animal Care and Use Committee (#707239) at the University of Iowa, in accordance with the National Institutes of Health Guide for the Care and Use of Laboratory Animals.

#### Mouse fixed-interval timing task

Mice were trained to perform an interval-timing task with a 12-sec interval[11]. Operant chambers (MedAssociates) were equipped with a nose poke hole with a yellow LED stimulus light (ENV-313W), a pellet dispenser (ENV-203-20), and a house light (ENV-315W). Behavioral chambers were housed in sound-attenuating chambers (MedAssociates). All behavioral responses including nose pokes and access to pellet receptacles were recorded with infra-red sensors. First, animals learned to make operant nose pokes to receive rewards (20-mg rodent purified pellets, F0071, BioServe). After fixed-ratio training, animals were trained in a 12-sec fixed-interval timing task in which rewards were delivered for responses after a 12-sec interval (Figure 1A). The house light was turned on to signal the start of the 12-sec interval. Early responses were not rewarded. Responses after 12 sec resulted in trial termination with reward delivery. Rewarded nose pokes were signaled by the house light turning off. Each trial was followed by a 30± 6-sec pseudorandom inter-trial interval that concluded with the house light turning on, signaling the beginning of the next trial. All sessions were 60 min long. Responses were summed into time-response histograms with 1-sec bins from 0 to 18 sec after trial start. For plotting, we used the MATLAB function ksdensity.m to estimate the probability density function time-response histograms with a bandwidth of 1, normalized to maximum response rate, and averaged across animals. We quantified timing using a measure of the curvature of time-response histograms. This metric is based on the cumulative distribution function’s deviation from a straight line; it is 0 when the time-response curve is flat during the interval but closer to 1 when more responses are at 12 s and time-response histograms are more curved. We and others have used this metric extensively to quantify timing because curvature is resistant to differences in overall response rate [11, 24-26]. Curvature indices and response rates were compared between the normal saline and the scopolamine groups using a two-tailed t-test; p<0.05 was interpreted as statistically significant.

**Figure 1.**
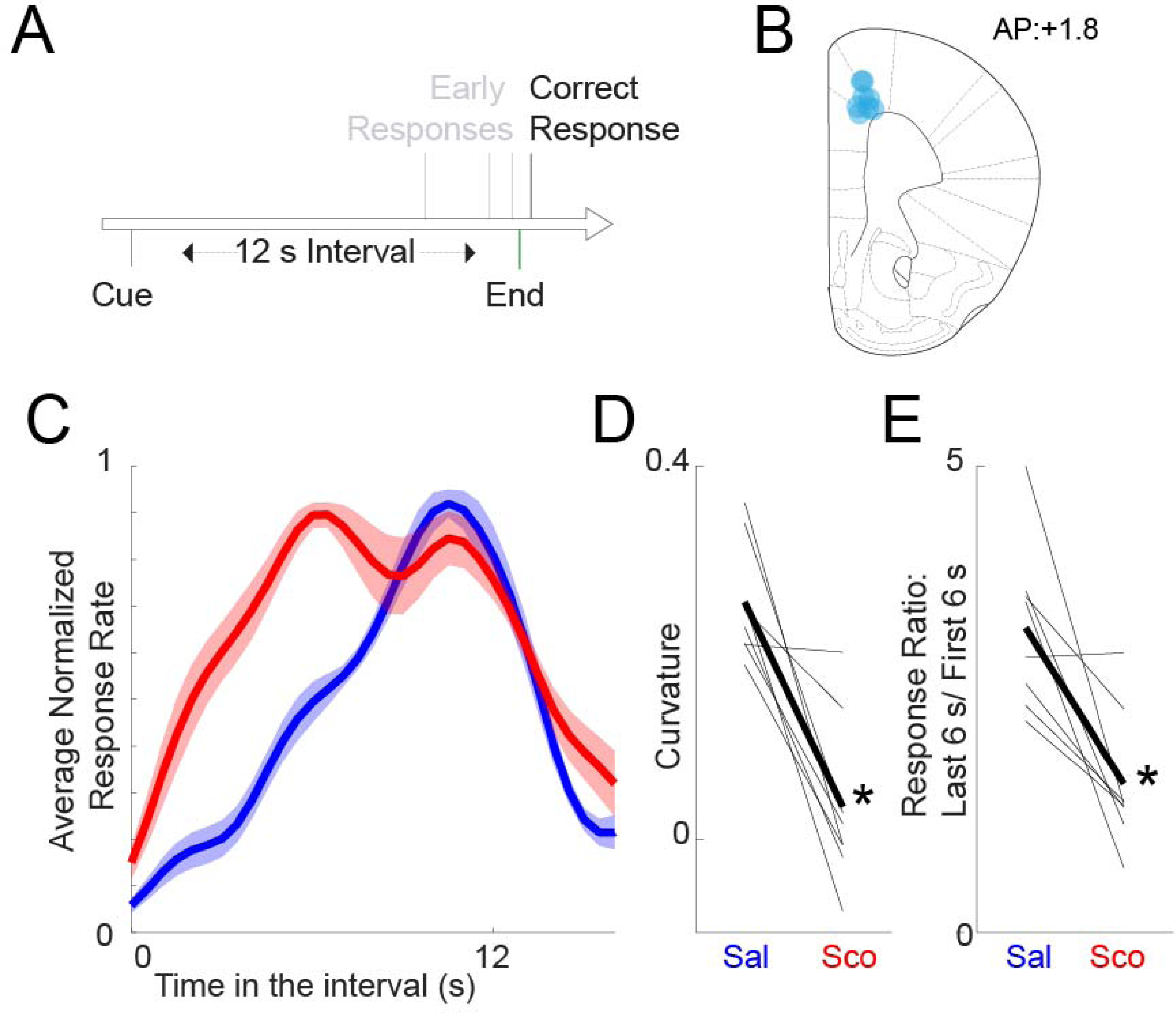
Scopolamine impairs interval timing. A) Interval timing task: Mice were trained to perform a fixed-interval timing task with a 12 s interval. The first response after 12 s led to a food reward; early responses were unreinforced. B) Recording electrode locations (blue dots) in the medial frontal cortex. C) Time-response histograms during fixed-interval timing. Data for mice administered saline are plotted in blue, and those for mice administered scopolamine are plotted in red. D) Timing can be quantified by computing a curvature index of time-response histograms; scopolamine decreases the curvature index, indicating flatter time-response histograms. E) We also noticed that animals with scopolamine responded more during the earlier portion of the interval. The ratio of responses in the last 6 vs. first 6 s closer to one, unlike in sessions with saline injected. Data from 8 mice; * =p<0.05 via paired t-test.

#### Surgical procedures

Mice trained in the 12-sec fixed-interval-timing task were implanted with recording microelectrode arrays (Microprobes) targeting the MFC prior to neurophysiology recordings. Briefly, mice were anesthetized using ketamine (100 mg/kg) and xylazine (10 mg/kg). A surgical level of anesthesia was maintained, with ketamine supplements (10 mg/kg) given hourly (or as needed) and regular monitoring for stable respiratory rate and absent toe pinch response. Mice were placed in the stereotactic equipment with non-rupturing ear bars. A heating pad was used to prevent hypothermia. Under aseptic surgical conditions, the skull was leveled between the bregma and lambda. A single craniotomy was drilled over the area above the MFC and three holes were drilled for skull screws. For recording experiments, animals were implanted (coordinates from the bregma: AP: +1.8, ML + 0.5, DV −1.8) with a microelectrode array configured as a 4×4 array of 50 μm stainless steel wires (200 μm between wires and rows; impedance measured *in vitro* at 400–600 kΩ; Microprobes). Electrode ground wires were wrapped around the skull screws. The electrode array was inserted while concurrently recording neuronal activity. The craniotomy was sealed with cyanoacrylate (“SloZap”, Pacer Technologies) accelerated by “ZipKicker” (Pacer Technologies), and methyl methacrylate (“dental cement”; AM Systems). Following implantation, animals were allowed to recover for two weeks before being reacclimatized to behavioral and recording procedures.

#### Neuronal ensemble recordings

Freely moving electrophysiological recordings were performed as described in detail previously [11, 24]. Following training in a 12-sec interval timing task and MFC implantation with 16-channel microelectrodes, mice were subjected to intraperitoneal injection with normal saline or scopolamine (1mg/kg, Sigma-Aldrich, S0929) while being recorded during the 12-sec interval timing task. Mice were connected to recording head stages and a cable without anesthesia. Neuronal ensemble recordings in the MFC were made using a multi-electrode recording system (Open Ephys). Raw wideband signal was high-pass filtered at 0.05Hz with total gain of 5000, and recorded with 16-bit digitization at 30k Hz sampling rate. To detect spikes, raw signals were rereferenced using common median referencing to minimize potential non-neural electrical noise, and band-pass filtered between 300 and 6000 Hz offline. Spikes were detected with a threshold of 5 median absolute deviations. A Plexon Offline Sorter was used to sort single units and to remove artifacts. PCA and waveform shape were used for spike sorting. Single units were identified as having 1) consistent waveform shape, 2) separable clusters in PCA space, 3) a consistent refractory period of at least 1 ms in inter-spike-interval histograms, and 4) consistent firing rates around behavioral events. Unique neurons were verified by constructing two-dimensional cumulative distribution probabilities from Pearson’s correlation coefficients of pair-wise waveform and inter-spike-interval comparisons, and using a one-tailed threshold of p<0.05. Spike activity was analyzed for all cells that fired at rates above 0.1 Hz. Local field potential (LFP) was recorded with bandpass filters between 0.05 and 1000 Hz. Statistical summaries were based on all recorded neurons. No subpopulations were selected or filtered out of the neuron database. Analysis of neuronal activity and quantitative analysis of basic firing properties were carried out with custom routines for MATLAB (all raw data and MATLAB scripts are available at our lab website: https://narayanan.lab.uiowa.edu/article/datasets). All behavioral events were recorded simultaneously using TTL inputs at 30k Hz. Peri-event rasters and average histograms were constructed around trial start.

We analyzed our neuronal data according to procedures described at length previously [11, 24]. For each neuron, we constructed peri-event spike data from −2 sec to 12 sec after trial start. As in the past, we defined time-related ramping neurons as those with a significant fit via linear regression of time vs. firing rate over the interval binned at 1 sec. Finally, we defined stimulus-modulated and response-modulated neurons as those with trial-by-trial changes in firing rate 0–200 msec after event onset compared to −500 to −300 msec prior to stimulus onset with a p<0.05 via a paired t-test. Pearson’s chi-squared test was used to compare the number of modulated neurons between saline and scopolamine sessions.

We analyzed neuronal patterns using PCA, which we have used to identify patterns of neuronal activity in an unbiased, data-driven manner [11, 27-29]. PCA was constructed from z-transformed peri-event time histograms over the entire interval binned at 0.1 sec and smoothed with a gaussian kernel over 5 bins. All neurons from 8 mice from sessions with saline or scopolamine infusions were included in PCA. We then used a t-test to compare PCs between saline and scopolamine sessions.

MFC LFP power was calculated in defined frequency bands (delta: 1-4 Hz, theta: 5-8 Hz; alpha: 9-12 Hz; beta: 13-30 Hz) during the interval (0-12 s) using wavelet-based time-frequency analyses.

#### Histology

When experiments were complete, mice were euthanized by injections of 100 mg/kg sodium pentobarbital. All mice were intracardially perfused with 4% paraformaldehyde. The brain was removed and post-fixed in paraformaldehyde overnight and immersed in 30% sucrose until the brains sank. Sections (40 µm) were made on a cryostat (Leica) and stored in cryoprotectant (50% PBS, 30% ethylene glycol, 20% glycerol) at −20°C, before being mounted onto slides with mounting media containing DAPI (Invitrogen P36962). Images were captured on an Olympus VS120 Microscope.

## Results

### Scopolamine impairs fixed-interval timing

Two past studies in rodents have demonstrated that scopolamine impairs interval timing [20, 21]. During neuronal recording sessions, we found that scopolamine infusion markedly changed time-response histograms during fixed-interval timing (Fig 1C). We calculated the *curvature* of time-response histograms to measure timing, a metric based on cumulative distribution functions that we and others have used in the past [11, 24-26]. We found that scopolamine significantly decreased the curvature of time-response histograms (0.25 +/- 0.02 vs. 0.03 +/-0.03 with scopolamine; paired t_(7)_=5.1, p=0.001; Fig 1D). Furthermore, we noticed that there were more responses early in the interval and that scopolamine decreased the ratio of responses late in the interval vs. early in the interval (last 6 sec divided by first 6 sec; 3.27 +/- 0.31 vs. 1.60 +/-0.26 with scopolamine, paired t_(7)_=3.9, p=0.006; Fig 1E). Scopolamine did not change the number of overall responses between 0 and 12 sec (86.3 +/-23.0 vs. 98.4 +/-28.5 with scopolamine) or the number of rewards (66.6 +/-5.3 vs. 63.8 +/- 5.3 with scopolamine). Taken together, our findings are consistent with past work demonstrating that scopolamine impairs interval timing [20, 21].

### Scopolamine does not change MFC ramping but increases stimulus-related processing

Prior work by our group and others has demonstrated that time-related ramping activity by MFC neurons is a key correlate of temporal processing [16, 23]. As scopolamine impairs interval timing, we hypothesized that this drug would impair time-related ramping. We tested this idea by identifying time-related ramping MFC neurons by linear regression (Fig 2A), and comparing the number of MFC ramping neurons in sessions with saline and scopolamine. In 8 mice, of 117 MFC neurons recorded during saline sessions, 41 exhibited time-related ramping (35%). Critically, a similar fraction of MFC neurons exhibited ramping with scopolamine (36 of 108, or 33%); this did not support our hypothesis.

**Figure 2.**
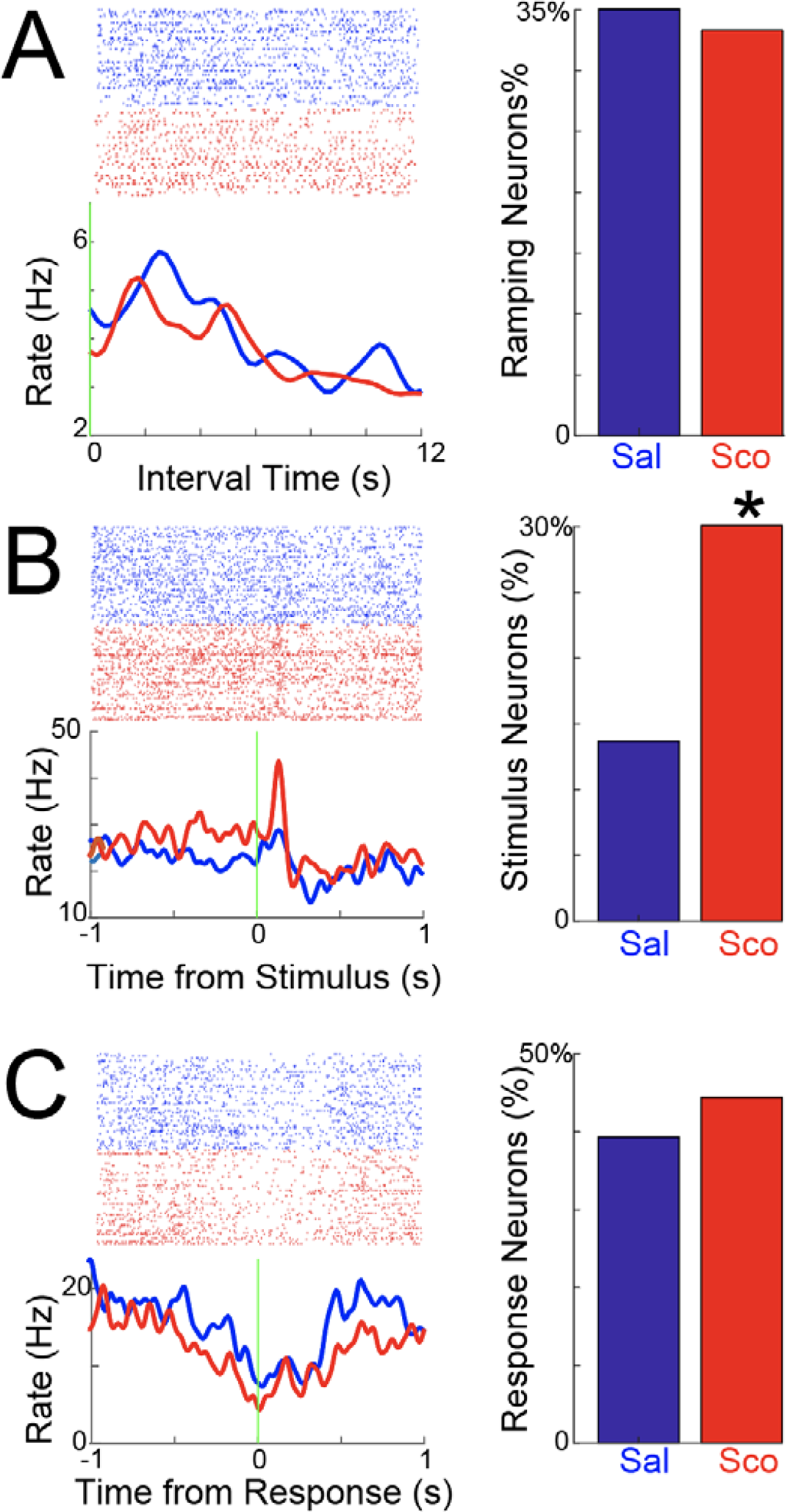
Scopolamine does not change MFC time-related ramping, but increases MFC stimulus-related activity. A) We identified MFC neurons with time-related ramping activity by linear regression. Crucially, scopolamine did not change the number of MFC ramping neurons. B) By contrast, scopolamine dramatically increased the number of MFC neurons with stimulus-related modulation. C) Scopolamine did not change MFC neurons with response-related modulation. Data from 8 mice; 117 MFC neurons recorded in saline sessions and 108 neurons recorded in scopolamine sessions; * =p<0.05 via χ^2^ test.

Next, we looked at other modulation patterns in MFC. Prefrontal regions can powerfully affect stimulus-processing [30-32]. During fixed-interval timing, this stimulus is a light that goes on at trial start (Fig 2B). Surprisingly, there were twice as many stimulus-modulated MFC neurons in scopolamine sessions (32 of 108, or 30%) compared to saline sessions (16 of 117, or 14%; X^2^=7.6, p=0.006). There were no differences in the number of response-modulated neurons (Figure 2C; 39% with saline vs. 44% with scopolamine). These data provide evidence that neither MFC ramping nor response-related activities are changed by scopolamine; by contrast, scopolamine increased MFC stimulus-related modulation.

When comparing average MFC neuronal ensemble activity in saline and scopolamine sessions, we noticed subtle differences late in the interval (increased activity at black arrow in Fig 3A vs. decreased activity at black arrow in Fig 3B). These resulted in different average activities of MFC neuronal ensembles with saline vs. scopolamine (Fig 3C). To capture these differences with data-driven techniques, we turned to principal component analysis, which have been used extensively to identify patterns in complex neuronal data[11, 27-29]. We found three common patterns. PC1, which explained 29% of variance, exhibited time-related ramping activity, consistent with extensive past work from our group (Fig 3D-E). Of note, this component did not change with scopolamine, consistent with our results above and contrary to our hypothesis. PC2, which explained 19% of variance, was broadly modulated across the interval, and also did not change with scopolamine. By contrast, PC3, which explained 12% of variance, had a more complex pattern, with a peak close to 7 sec in the interval. Of note, in the scopolamine sessions there was a more negative score with PC3 compared to saline sessions (Fig 3F, PC3; t_(223)_=2.4, p<0.02). These data provide further evidence that scopolamine did not change MFC ramping but could change more complex aspects of MFC neuronal ensemble activity.

**Figure 3.**
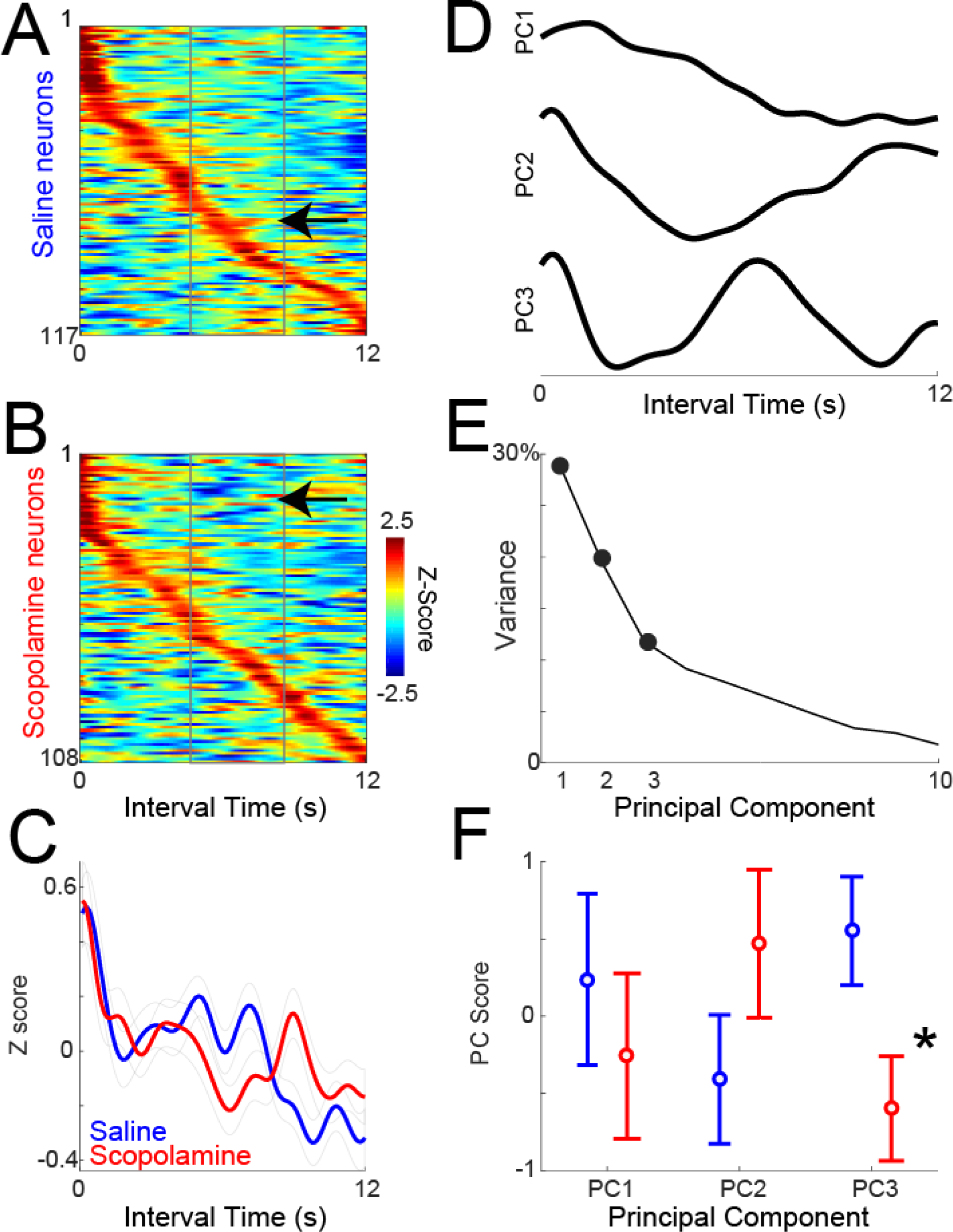
Neuronal ensemble effects of scopolamine: A) Average Z-scored neuronal activity during the interval shown for all MFC neurons treated with saline, and B) scopolamine, sorted by peak activity. We noticed subtle differences in activity, with more activity in saline (arrow in A), vs. less activity in scopolamine (arrow in B) late in the interval. Activity binned at 10 ms and smoothed over 5 bins. C) There were differences during the interval between average MFC neuronal activities in saline vs. scopolamine sessions. D) To quantify these differences using data-driven techniques, we turned to principal component analysis, which identified 3 major patterns. PC1 had ramping patterns, PC2 was modulated during the interval, and PC3 had more complex modulation. E) Fraction of variance explained by each component. F) Only PC3 was different between saline and scopolamine sessions. Data from 8 mice; 117 MFC neurons recorded in saline sessions and 108 neurons recorded in scopolamine sessions; * =p<0.05 via t-test.

Finally, we examined MFC LFP oscillatory power during the interval (0-12 s; Figure 4). We found no consistent changes in delta (1-4 Hz), theta (5-8 Hz), alpha (9-12 Hz), or beta activity (13-30 Hz). In summary, our results suggest that scopolamine impaired interval timing and enhances stimulus-related processing in MFC without changing MFC temporal processing or MFC LFPs.

**Figure 4.**
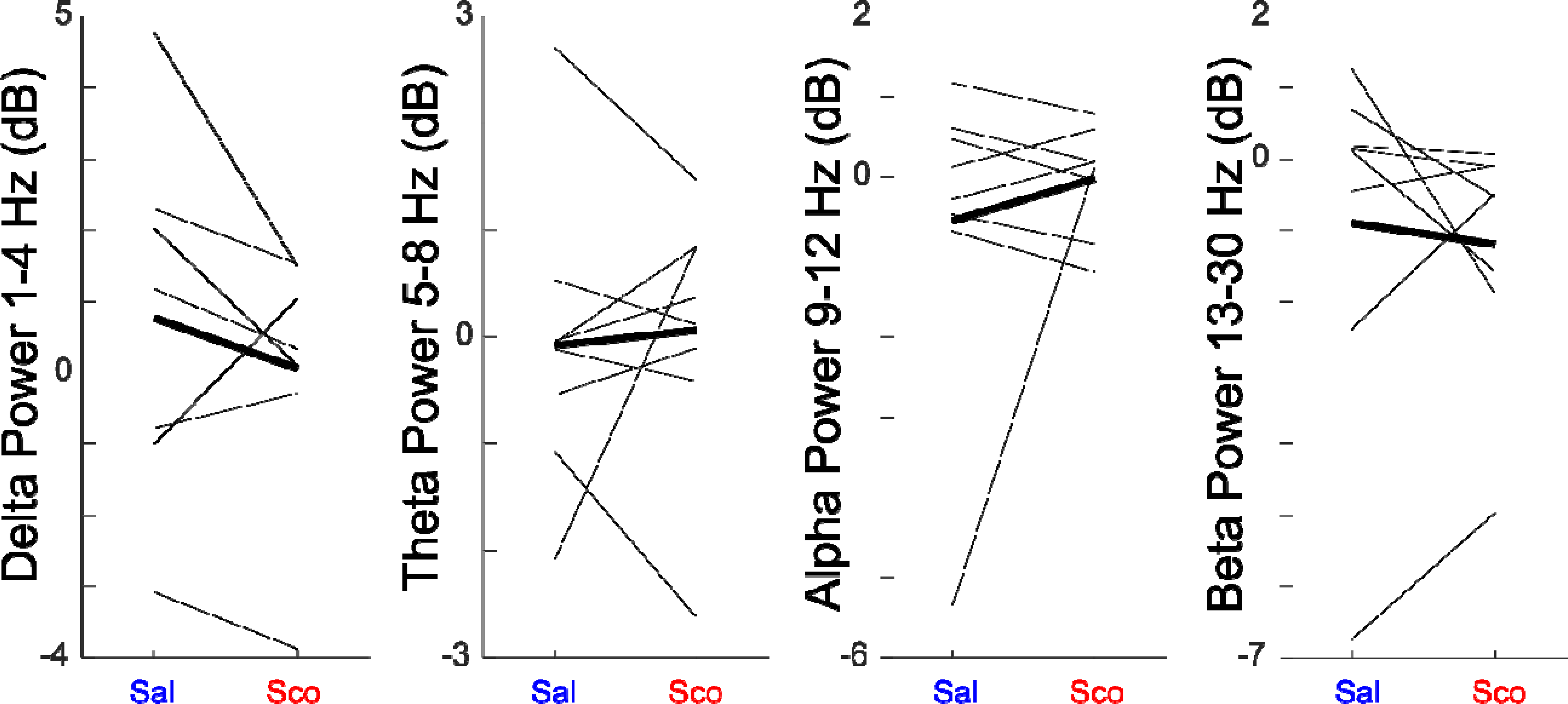
Scopolamine does not change MFC LFP activity: We measured MFC LFPs from 8 mice in delta, theta, alpha, and beta power in saline and scopolamine sessions. We did not observe consistent differences between saline and scopolamine sessions. Data from 8 mice.

## Discussion

We tested the hypothesis that scopolamine would attenuate time-related ramping activity in the MFC. We found no evidence that scopolamine changed MFC ramping by linear regression or PCA. To our surprise, we found that stimulus-related processing was increased in the MFC with scopolamine. These data provide insight into how scopolamine might influence cortical circuits during interval timing.

Many cognitive behaviors are impaired by the muscarinic cholinergic inhibitor scopolamine, often through attentional and stimulus-processing deficits [33, 34]. Scopolamine reliably causes timing deficits in rodents [20, 21], and here we report similar deficits during fixed-interval timing. Surprisingly, MFC ramping is intact with scopolamine administration. There are two possibilities that might account for this result. First, we and others have identified neuronal ramping as a key mechanism of temporal processing in the MFC [16, 22, 23, 35], but it is possible that ramping is not explicitly related to timing. In this case, others have proposed temporal computations based on oscillatory activity [36], and we note that PC3 has oscillatory features, although the period appeared to be longer than 1 second. Scopolamine may affect neuronal oscillatory activity that was not detected by our analyses. A second possibility is that scopolamine affects stimulus-processing mechanisms in the MFC and this triggers animals to respond despite intact MFC temporal processing [31, 37]. The increase in responses early in the interval is supportive of this idea.

Previous studies have shown that acetylcholine is crucial for stimulus processing. Microdialysis has revealed that acetylcholine is increased during tasks involving sustained attention, changes in ambient light, anticipation of a meal, motor activity, and handling, while amperometry has indicated that MFC acetylcholine can increase for reward-predictive cues [38-43]. Acetylcholine can also be increased during attentionally demanding tasks such as 5-choice serial reaction-time tasks [31, 44]. Lesioning of MFC cholinergic projections impairs the processing of fast but not slow stimuli, as well as causing marked deficits in visual attentional performance [45]. To our knowledge MFC cholinergic projections have never been studied in interval timing, although cholinergic projections to the visual cortex affect learning of temporal intervals but not interval-timing performance [46]. Lesioning of MFC cholinergic projections decreased stimulus-related activity and the performance of a visual attention task, contrary to our results here with scopolamine, a muscarinic antagonist [47]. In primates, cholinergic deafferentation can affect working memory, and cholinergic effects on working memory and attention occur as a result of the direct effects on stimulus processing [48, 49]. These studies broadly support a role for cortical cholinergic circuits in task-related stimulus processing.

Our results here are quite different from those for systemic or focal manipulations of dopaminergic circuits in the frontal cortex or striatum [12, 27, 50, 51]. While dopaminergic manipulations also reliably affect interval timing, our work indicates that manipulating prefrontal dopamine via D1-type dopamine receptors affects both time-related ramping activity and ∽4 Hz rhythms [11, 27]. Scopolamine appears to have distinct effects on MFC circuits enhancing stimulus-related activity while leaving MFC ramping and MFC LFPs unchanged. These data suggest that cholinergic and dopaminergic manipulations have distinct and specific effects on cortical circuits. Future studies with cell-type specific resolution might be able to further resolve themes of cortical cholinergic vs. dopaminergic circuits.

Patients with AD and PD have deficits in timing [10, 11, 15, 52-54]. We are not aware of studies of interval timing in DLB patients. PD and AD have prominent cholinergic deficits [1, 2]. Of note, cholinergic drugs can improve cognition in patients with AD, PD and DLB [55]. However, little is known about the relevant cholinergic circuit mechanisms of these effects.

Our study has several limitations. First, scopolamine was administered systemically and is a poor model of cognitive dysfunction in dementia, as it can act on all muscarinic acetylcholine receptors in the brain [33] and other brain systems[56]. Furthermore, it can have non-specific locomotor and autonomic effects, although we did not observe an increased response rate and observed highly specific neuronal effects on stimulus-processing in this study. Cholinergic interneurons in the striatum or the MFC may also be critical mediators of cognitive processing [57]. Finally, we are unsure if scopolamine directly modulates MFC stimulus-related neurons or modulates other brain areas. Nevertheless, our results lay important groundwork for highly specific investigation of cholinergic circuits using cell-type specific methods in future work.

In conclusion, we performed neuronal ensemble recording from the MFC of freely moving rodents, and found that muscarinic cholinergic inhibition caused timing deficits and hyper-stimulus-modulation during interval timing. MFC temporal processing was not significantly affected. These results are consistent with the consensus that acetylcholine plays major roles in attention, working memory and stimulus processing, while the dopaminergic system is crucial for the neuronal “ramping” activities and an internal pacemaker. Neuromodulation therapies targeting the cholinergic circuits have potential as treatments for AD as well as DLB [55]. Data from this study will help us understand the cholinergic circuits and may have relevance for diseases involving cholinergic deficits, such as AD and DLB.

## Contributors

QZ, YK and NN designed the study. QZ, DJ, and TL performed the experiments and collected the data. QZ, YK and NN analyzed the data. QZ, YK and NN interpreted the results and wrote the manuscript.

## Acknowledgements

NN is supported by R01MH116043A1. QZ is supported by the NINDS R25 grant, a pilot project grant from the Aging Mind and Brain Initiative at University of Iowa, and the physician scientist training program at University of Iowa. QZ is a trainee of the University of Iowa Clinical Neuroscientist Training Program (CNS-TP).

## Conflicts of interests

There are no conflicts of interests.

